# RePaRank: An Efficient Architecture for Antibody-Antigen Interface Prediction by Proximity Ranking

**DOI:** 10.64898/2026.03.03.708462

**Authors:** Jakub Bednarek, Bartosz Janusz, Konrad Krawczyk

## Abstract

The prediction of protein-protein interactions is central to structural biology, yet leading models are often computationally expensive, creating an accessibility gap for many high-throughput applications. Furthermore, common evaluation metrics such as binary contact prediction can be unreliable. In this work, we address both challenges. We introduce RePaRank, a computationally efficient deep learning architecture with 39.4 million parameters that predicts antibody-antigen interfaces by reframing the problem as a proximity ranking task in a learned embedding space. We also propose the Precision AUC, a robust, ranking-based metric that provides a more stable assessment of model performance than traditional binary methods. Our experiments show that RePaRank consistently outperforms benchmark models in paratope prediction and is highly competitive in epitope prediction among models that do not require external resources such as Multiple Sequence Alignments (MSA). RePaRank offers a practical and powerful tool for the immunoinformatics community.

## 1. Introduction

The application of machine learning to structural biology has revolutionized the field, culminating in models such as **AlphaFold2** [1] and **RoseTTAFold** [2] that have largely solved the single-chain protein folding problem with unprecedented accuracy. The next frontier is the prediction of protein-protein interactions (PPIs), the molecular basis of nearly every biological process. Predicting the interface between an antibody and an antigen is a particularly critical challenge in immunology and therapeutic development. While state-of-the-art models have demonstrated remarkable capabilities, their success often depends on deep Multiple Sequence Alignments (MSAs) and massive computational resources, creating an accessibility gap for many researchers and limiting their use in high-throughput screening.

To bridge this gap, a new generation of MSA-free models has emerged, often leveraging large protein language models such as **ESMFold** [3]. While powerful, these models can still be computationally demanding. We propose **RePaRank**, a computationally lean, MSA-free deep learning architecture for antibody-antigen interface prediction with only **39.4M** of parameters, while competing models range from **300M to 3.5B** parameters. RePaRank is designed for predicting antibody-antigen interfaces by reframing the problem from a binary classification of atomic contacts to a **proximity ranking task**, where the model learns to organize residue representations in N-dimensional space based on the relationships between them in the protein structure, repelling distant pairs of residues and attracting close ones. The problem formulation is sufficiently general to facilitate downstream applications in established domains, such as contact map prediction.

A parallel challenge in the field is the evaluation of model performance. For machine learning predictions to be practically adopted by biologists, they must achieve high precision to minimize the significant cost and effort of experimental validation. However, common metrics such as binary contact prediction, that rely on arbitrary distance cutoffs, can be misleading. Even more comprehensive methods such as the Precision-Recall curve have notable drawbacks; their results are highly dependent on the class distribution of the test data, and they do not produce a single, easily comparable numeric score to rank different models [4]. To address these limitations, we propose the Precision AUC, a robust, ranking-based metric that provides a more stable and informative assessment of a model’s ability to prioritize true interface residues across all decision thresholds.

Our experiments demonstrate that RePaRank, evaluated with this improved metric, consistently outperforms benchmark models in paratope prediction and is highly competitive in epitope prediction. By combining an efficient architecture with a more meaningful evaluation framework, RePaRank offers a practical, accessible, and powerful tool for the immunoinformatics community.

## 2. Materials and Methods

This section details the framework for developing and validating RePaRank, our model for identifying the binding interfaces of antibody-antigen complexes. We describe the data preparation and benchmarking procedures, the model architecture and its components, and the specific training strategy employed to ensure the reproducibility of our experiments.

### 2.1 Data

The datasets for this study were derived from NAStructural Database (APE; [5]). We employed two distinct interface subsets: antibody-antigen (Ab-Ag) and general protein-protein (PP). The Ab-Ag subset serves as our primary dataset for the central task of epitope and paratope identification, while PP subset was used exclusively at the model pre-training stage in one of our experiments.

To ensure a rigorous and unbiased evaluation, we designed a splitting protocol to generate non-redundant training and test sets, thereby mitigating data leakage. Our methodology combines a temporal split with sequence identity clustering. The test set is composed exclusively of structures published after a cutoff data of September 30, 2021. This cutoff was selected to align with the training data of our benchmark models (Chai-1 [6], Boltz-1 [7], and ESMFold [3]), ensuring a fair comparison.

To enforce structural and sequence novelty in the test set, we partitioned the data based on sequence identity. A threshold of 80% sequence identity was applied for the highly conserved antibody chains, while a stricter 30% threshold was used for antigens (in the Ab-Ag set) and all protein chains within the PP subset. A complex was assigned to the test set only if it met two criteria: its publication date was after the specified cutoff, and all of its protein chains were sufficiently dissimilar (per the thresholds) from any chain in the training set. All remaining complexes were allocated to the training set.

The final distribution of the datasets across the training, validation, and test splits is summarized by the number of partitions and their constituent original complexes. The Ab-Ag dataset comprises 727 training (1850 complexes), 73 validation (167 complexes), and 297 test partitions (585 complexes). Similarly, the PP dataset contains 3299 training (36535 complexes), 348 validation (2630 complexes), and 466 test partitions (1406 complexes).

### 2.2 Architecture

The **RePaRank** architecture is designed to address the specific task of identifying protein interactions sites, such as epitopes and paratopes, rather than generating full 3D structures. We reformulate this challenge as a ranking problem focused on determining the closest inter-residue interactions within a protein complex. The core functionality of our model is to provide a ranking for all possible residue-residue pairs, where the ranking reflects their spatial proximity in the native 3D structure. This approach is highly versatile; for instance, the top-ranked pairs between an antibody and an antigen define the binding interface, while top-ranked pairs between a heavy and light chain can identify intra-antibody contacts.

By reformulating the problem from 3D coordinate prediction to proximity ranking, our method bypasses the need to explicitly model the complex geometric and physical constraints governing protein folding. Instead, the model learns a simpler objective: to project residues into an N-dimensional embedding space where the Euclidean distance between any two embeddings correlates with their actual spatial proximity in the ground truth structure. We show in Figure 1 heatmaps depicting pairwise distances between heavy and antigen chains for ground-truth 3D coordinates and predicted N-dimensional embeddings.

**Figure 1.**
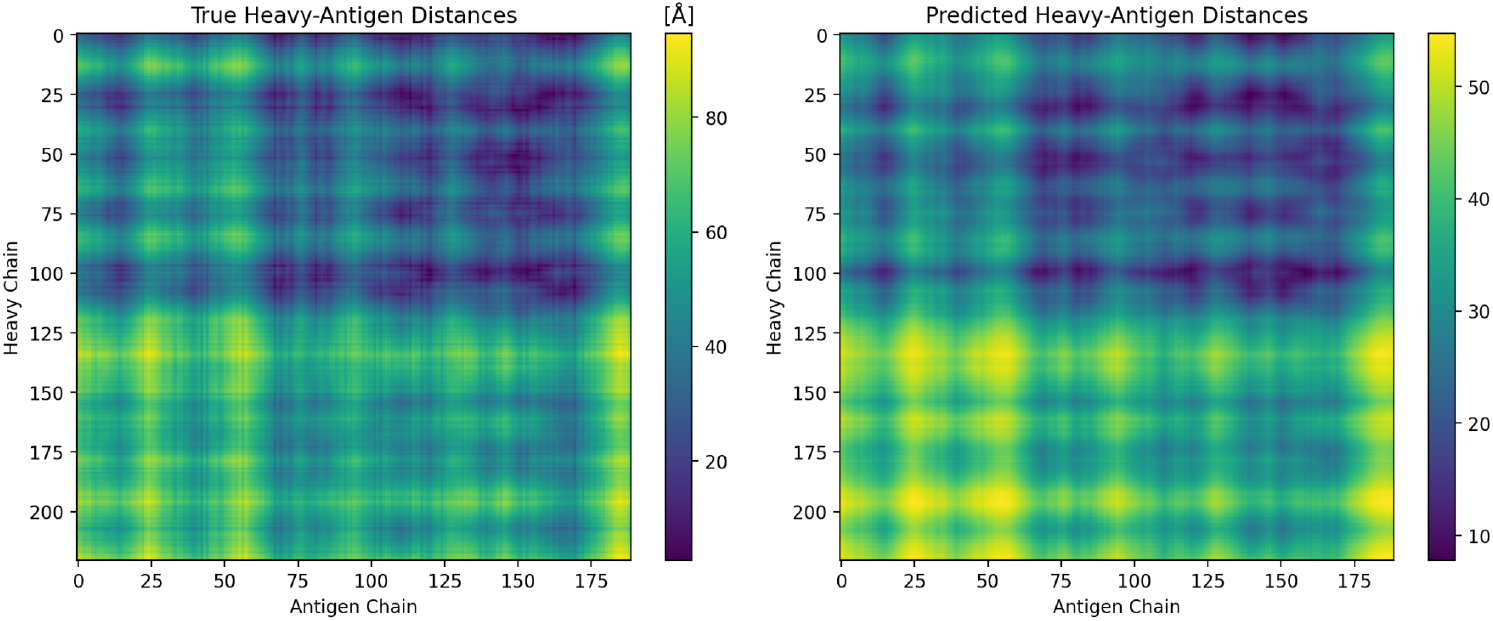
Comparison of the ground truth pairwise distances (left) with the distances predicted by RePaRank (right) for an example antibody heavy chain (y-axis) and antigen chain (x-axis) from *7dk2* PDB example. The example was placed in the test dataset. The ground truth map is based on the minimum heavy-atom Euclidean distance in the experimental structure and is expressed in units of Angstroms (Å), while the predicted map is derived from the Euclidean distance between residue embeddings in the learned 512-dimensional space. A strong visual correspondence demonstrates the model’s ability to accurately represent the complex’s true spatial relationships in its embedding space. Note that the absolute values on the color bars differ, as one represents physical distance (in Ångströms) and the other represents distance in an abstract, learned space.

### 2.3 Components

The proposed architecture consists of two main modules: Sequence Encoder and Complex Encoder:

- **Sequence Encoder:** This module processes each input protein chain independently to generate initial, context-independent residue embeddings. An embedding layer first transforms the one-hot encoded amino acid sequence into dense vectors. These are combined with positional encodings and passed through three convolutional residual layers to capture local features. Finally, a stack of eight self-attention layers processes the sequence to produce representations that are contextualized within each individual chain. The module, optionally, accepts structures in the form of residue locations (3D coordinates of backbone atoms; context-independent).
- **Complex Encoder:** The latent representations from all chains in the complex are concatenated and passed to this module. Complex Encoder processes these representations jointly, allowing the model to learn inter-chain dependencies. It consists of eight self-attention layers that model the entire complex, outputting the final, context-aware N-dimensional embedding for each residue.

The final output is a 512-dimensional embedding for each residue. All self-attention layers utilize a multi-head attention mechanism with 16 heads and a feed-forward network with a hidden dimension of 1024. For regularization, dropout with a rate of 0.15 is applied within the feed-forward layers. The RePaRank architecture and data processing pipeline, including the training stage, are shown in Figure 2.

**Figure 2.**
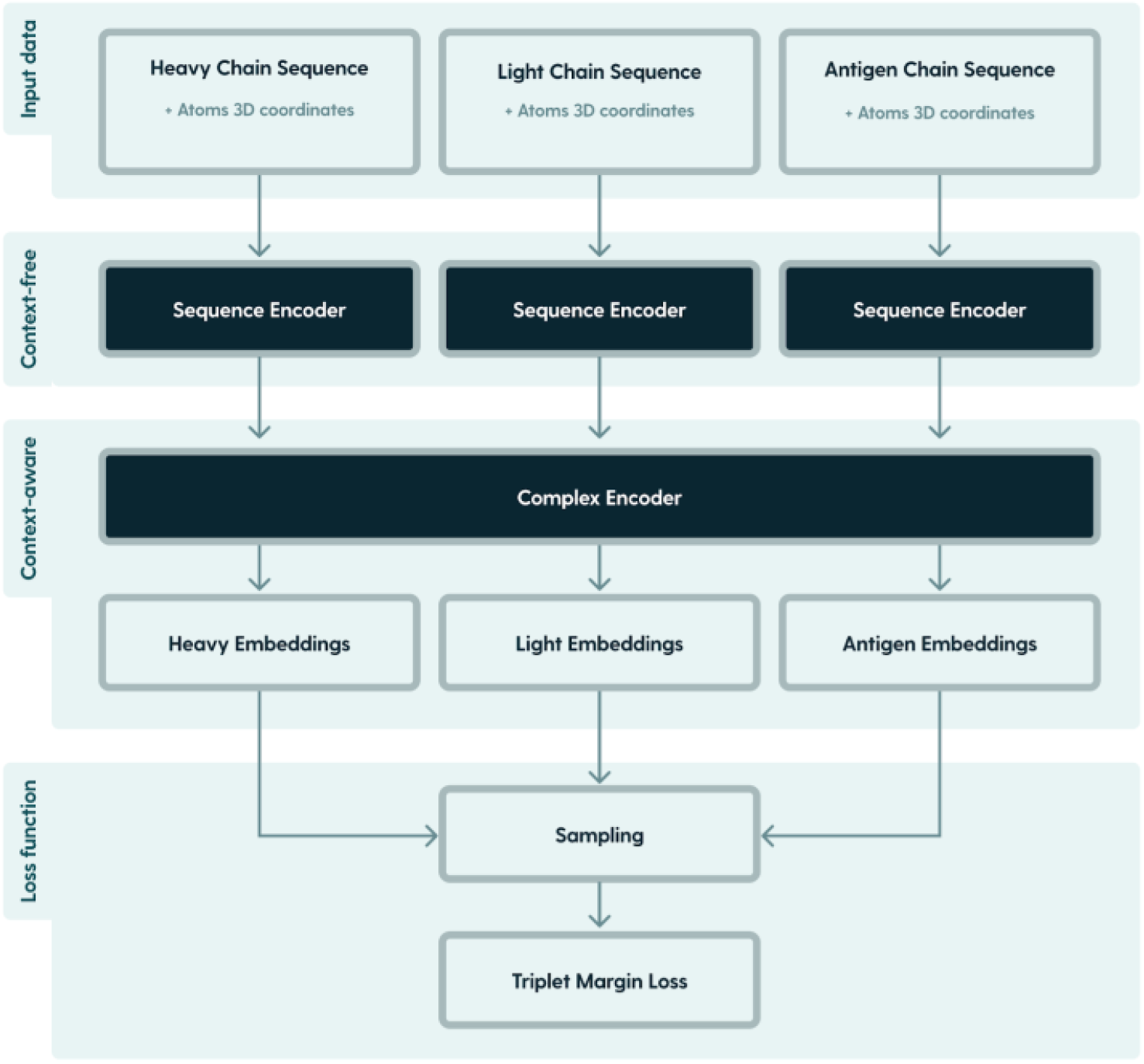
The RePaRank architecture. The model employs a two-stage encoding process. First, a Sequence Encoder processes each input protein chain (with optional structural data) independently to generate context-free residue embeddings. These initial embeddings are then passed to a single Complex Encoder, which processes all chains jointly to produce final, context-aware embeddings that capture inter-chain dependencies. During training, these embeddings are optimized using a Triplet Margin Loss that is informed by an anchor-positive-negative sampling strategy.

The complete architecture has **39.4 million** trainable parameters. This model size provides sufficient capacity to learn complex inter-residue dependencies while remaining computationally manageable for training and inference on a single commercially available GPU.

During inference, the model generates 512-dimensional embeddings for all residues in a given complex. By computing the pairwise Euclidean distances between these embeddings, we generate a ranking of the most probable residue-residue interactions, where smaller distances imply a higher likelihood of contact.

### 2.4 Training

We trained the model to map 3D spatial relationships into the N-dimensional embedding space using a triplet margin loss function [8]. During training, we sample triplets of residues *(a,p,n)*- representing an **anchor**, a **positive**, and a **negative** example - from the training complexes. These triplets are not restricted to residues from the same chain.

A triplet is defined based on the ground truth 3D structure, where the distance between the anchor and positive *d(a, p)*is less than the distance between the anchor and the negative *d(a, n)*.We define the ground truth distance *d* as the minimum Euclidean distance between any two heavy atoms of the respective residues.

The objective of the triplet loss is to train the model’s embedding function *f(*.)to satisfy the same distance relationship in the learned N-dimensional space. Specifically, the loss function encourages the distance between the embeddings of the anchor and positive *D(f(a), f(p))*to be smaller than the distance between the embeddings of the anchor and negative *D(f(a), f(n))*by at least a margin *m*:

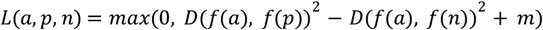

Here, *D* represents the Euclidean distance in the N-dimensional space. This training scheme effectively teaches the model to arrange residue embeddings such that their relative distances reflect the geometry of the original protein complex; *m* was set to 1.0.

The architecture was trained with the AdamW [9] optimization method with a learning rate 5e-5 and all other hyperparameters set to default values. The learning rate was reduced by the factor 0.1 every stagnation stage (no improvement on epitope identification). The training was performed for 500 epochs. The total training time was approximately 2 hours on a single machine with NVIDIA GeForce RTX 3090.

### 2.5. Availability

The datasets used for processing and model checkpoints generated during this study are publicly available to ensure reproducibility.

#### Model Checkpoints

We provide four distinct model checkpoints corresponding to different input modalities and pre-training stages:

- Seq: Sequence-only input.
- Seq + Struct: Sequence with structural features.
- Seq (PP pretrained): Sequence-only model with protein-protein pre-training.
- Seq + Struct (PP pretrained): Structural model with protein-protein pre-training.

All model artifacts are hosted on Google Drive and can be accessed at: [Google Drive](https://drive.google.com/drive/folders/1LNei4WY8SjG47dnUFKuFRxZLpd3IOQy9).

#### Datasets

The antibody-antigen and protein-protein interface subsets used for training and evaluation are available via the NA-Structural Database at: NaturalAntibody.com.

## 3. Results

### 3.1. Evaluation Methodology

We quantitatively assess model performance using a precision-based methodology inspired by information retrieval. This framework allows for a direct comparison between our model’s predictions and ground truth structures, as well as with other structural modeling methods.

For any given protein complex, our evaluation involves three key steps:

1. **Generate Interaction Rankings:** We create two ranked lists of all possible residue-residue pairs. The **ground truth ranking** is based on the true spatial proximity, where pairs with the smallest minimum heavy-atom distance rank highest. The **predicted ranking** is based on the Euclidean distance between the model’s residue embeddings, where pairs with the smallest embedding distance rank highest.
2. **Calculate Precision at Quantiles:** We evaluate the prediction quality by calculating precision at various retrieval depths. We define *N* quantiles, ranging from 0 to 1. For each quantile q ∈ [0,1], we consider the top *q*-fraction on residue pairs from the predicted ranking. We then calculate the precision for this set, which is the proportion of pairs that are also present in the top *q*-fraction of the ground truth ranking.
3. **Compute Precision AUC:** To aggregate the performance into a single, robust metric, we plot precision as a function of the quantile and calculate the **Area Under the Curve (AUC)**. This **Precision AUC** score serves as our primary evaluation metric.

The Precision AUC provides an intuitive measure of performance. A score of 1.0 indicates a perfect prediction, where the model’s ranking of residue proximities exactly matches the ground truth. A score of 0.5 corresponds to a random prediction, serving as a critical baseline.

A key advantage of this methodology is its versatility. It can be applied to assess any subset of interactions, such as those within a single chain (intra-chain) or between different chains (inter-chain), enabling detailed analysis of a model’s ability to capture specific types of protein contacts. This standardized metric ensures a fair and rigorous comparison across different predictive models.

### 3.2. Experiments

#### Experimental Setup

To validate our proposed architecture, **RePaRank**, we conducted a series of experiments to evaluate its performance against several reference, representative, models on our curated test set.

We benchmarked RePaRank against three structural prediction models: **Chai-1, Boltz-1**, and **ESMFold**. For our own model, we evaluated four distinct configurations to assess the contribution of different input features:

– **RePaRank (Seq):** The model was provided with only amino acid sequences.
– **RePaRank (Seq+Struct):** Sequences were independently^1^ supplemented with 3D structures.
– **RePaRank: (Seq PP pretrained):** The model with only amino acid sequences, pretrained on PP dataset.
– **RePaRank: (Seq+Struct PP pretrained):** The model accepting sequences with their independent structures, pretrained on PP dataset.

The primary metric for our main evaluation was the Precision AUC, calculated for an epitope (minimum distances of an antigen’s residues to an antibody) and a paratope (minimum distances of an antibody to an antigen, separately for heavy and light chains).

#### Epitope/Paratope prediction performance

We evaluated the models on their ability to call paratope/epitope residues without predicting the pairwise contacts between the two. We create a list of all pairs of residues between two chains and sort them based on their distances (for ground-truth, the true distance; for predicted, the distances between vectors). For the resulting rankings, we calculate the Precision AUC to check how well predicted ranking reproduces ground-truth. The Precision AUC metric is sensitive to the quality of epitope and paratope prediction (since these are the highest ranked positions) while also measuring the quality of prediction for residues that directly and indirectly determine epitope and paratope locations. The performance of models is summarized in Table 1. The results demonstrate that our model, RePaRank, outperforms all benchmark models in paratope prediction. Notably, the benchmark models Chai-1 and Boltz-1 achieve their best results when leveraging Multiple Sequence Alignments (MSA), a process that requires searching massive external sequence databases during inference. When compared to these models in a setting without access to external data (i.e. without MSA), RePaRank outperforms other models in all measured metrics.. Additionally, to address the limited training data for antibody-antigen complexes, we explored transfer learning. This involved pre-training RePaRank on the general PP dataset before fine-tuning it on the primary Ab-Ag dataset. This strategy was tested for both the sequence-only and the sequence-with-structure configurations or our model.

**Table 1.**
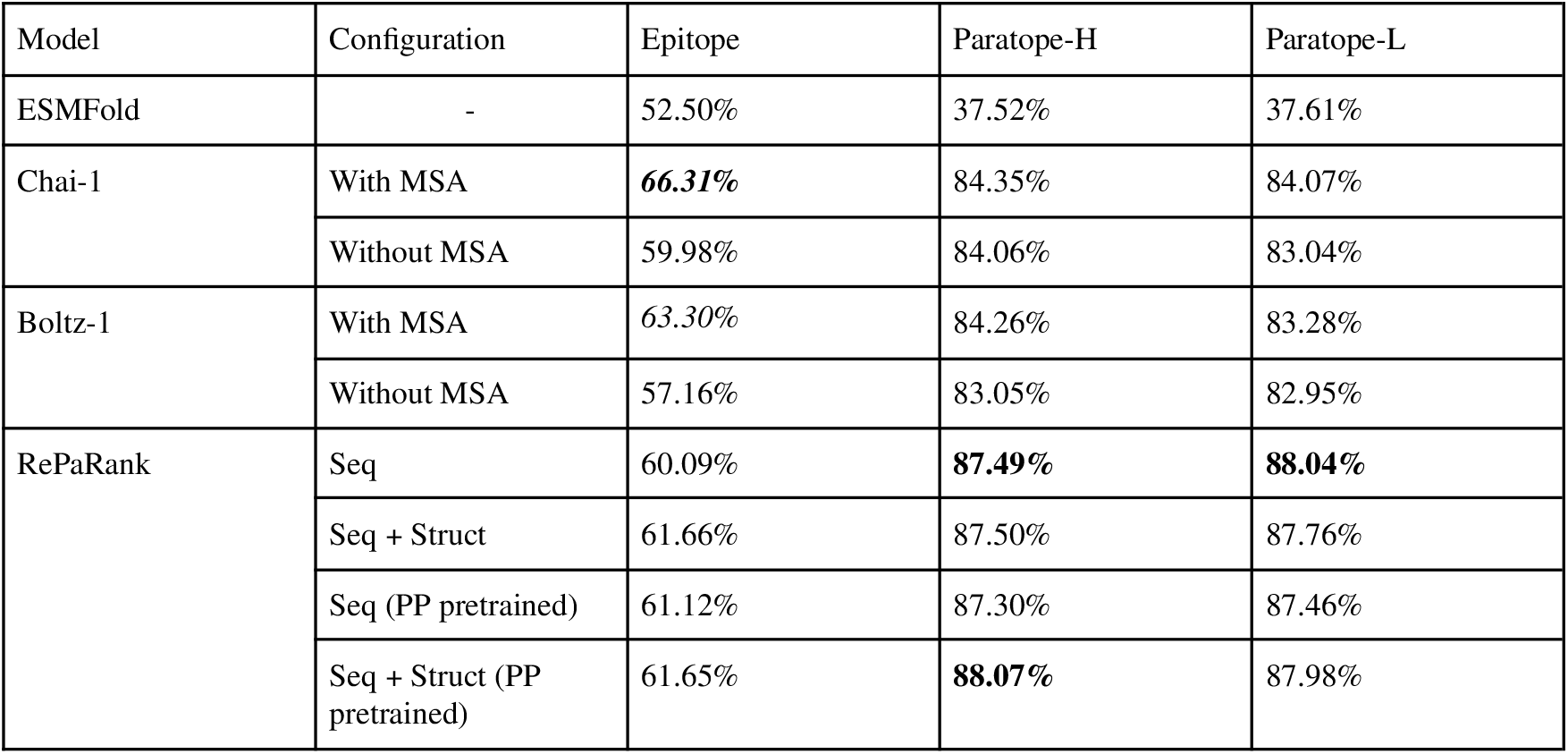
Comparative Performance on Interface Prediction. Precision AUC (%) scores for epitope and paratope identification across all models and configurations. While Chai-1 with MSA achieves the best epitope score, RePaRank consistently outperforms all competitors on both heavy and light chain paratope prediction, even in its simplest sequence-only configuration. A Precision AUC value of **50%** serves as the baseline for a non-informative model, representing the expected performance of a purely random method.

| Model | Configuration | Epitope | Paratope-H | Paratope-L |
| --- | --- | --- | --- | --- |
| ESMFold | - | 52.50% | 37.52% | 37.61% |
| Chai-1 | With MSA | **66.31%** | 84.35% | 84.07% |
|  | Without MSA | 59.98% | 84.06% | 83.04% |
| Boltz-1 | With MSA | 63.30% | 84.26% | 83.28% |
|  | Without MSA | 57.16% | 83.05% | 82.95% |
| RePaRank | Seq | 60.09% | **87.49%** | **88.04%** |
|  | Seq + Struct | 61.66% | 87.50% | 87.76% |
|  | Seq (PP pretrained) | 61.12% | 87.30% | 87.46% |
|  | Seq + Struct (PP pretrained) | 61.65% | **88.07%** | 87.98% |

To visualize the trade-off between model complexity and predictive power, we show Epitope (Figure 3) and Paratope (Figure 4; for a heavy-antigen interface) Precision AUC against the number of parameters for each model. This analysis highlights our model’s efficiency in achieving high performance without an excessively large parameter count.

**Figure 3.**
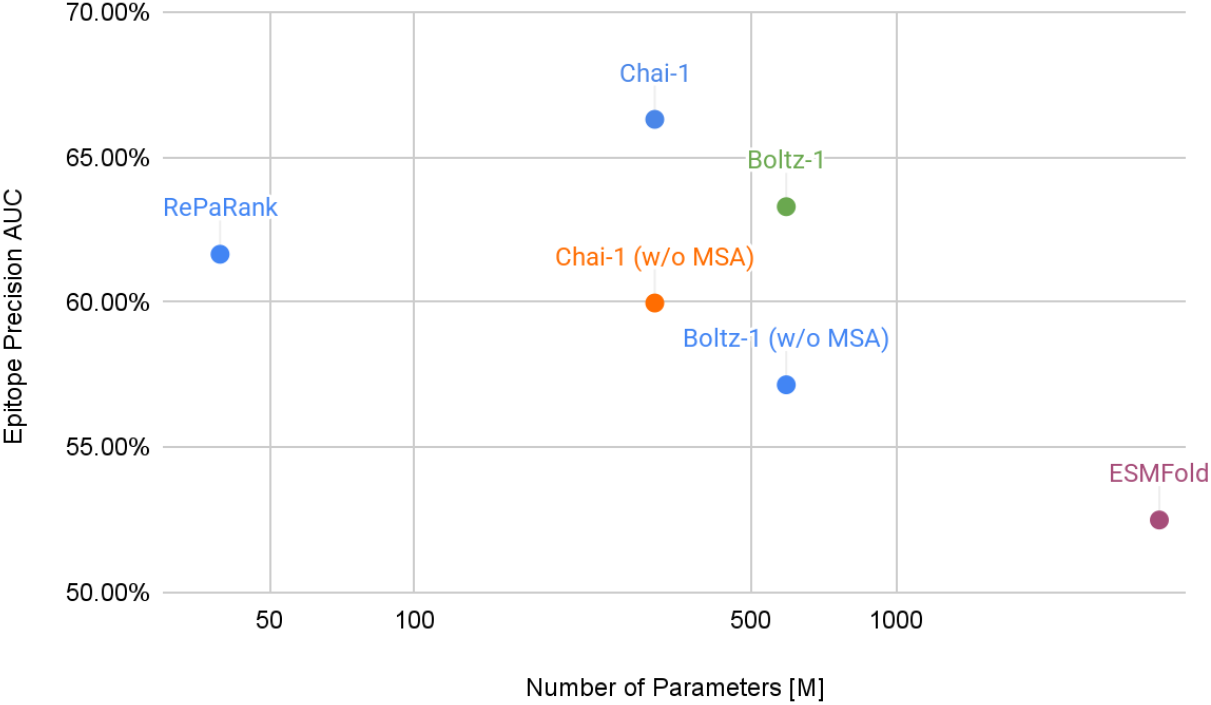
Performance vs. Model Complexity. The plot compares Epitope Precision AUC against the number of trainable parameters (log scale). RePaRank achieves the highest performance among all self-contained models that do not rely on an external Multiple Sequence Alignment (MSA) database, demonstrating a superior trade-off between model size and predictive accuracy.

**Figure 4.**
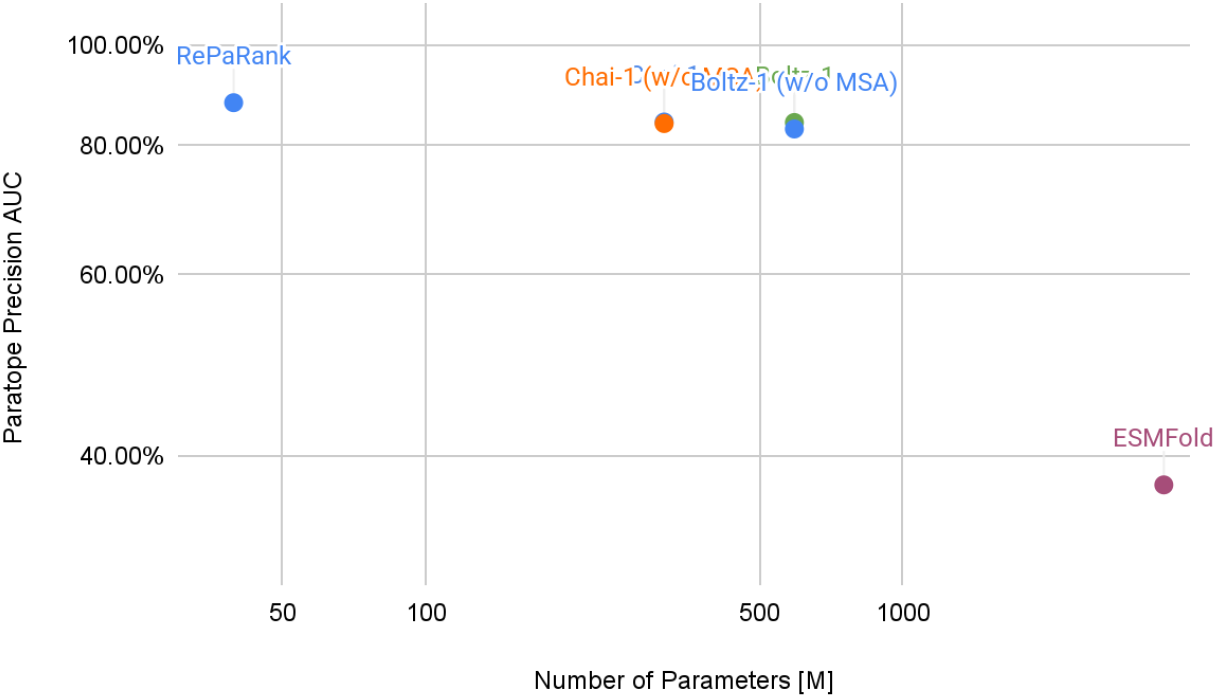
Performance vs. Model Complexity. RePaRank achieves the highest Paratope Precision AUC with substantially fewer parameters than competing architectures. Notably, it surpasses models utilizing MSA-based protocols, demonstrating superior parameter efficiency without the need for external evolutionary data.

#### Pairwise Contact Prediction

We evaluated the model on a binary contact prediction task. For the ground truth a contact was defined as any residue pair with a minimum heavy-atom Euclidean distance of less than 4.5Å. Although RePaRank operates in N-dimensional embedding space rather than 3D coordinates, it is trained so the smaller embedding distances directly correspond to closer physical proximities. We leveraged this property by finding a distance threshold in the embedding space that optimally maps to the 4.5Å physical contact definition. We propose a simple method used on top of RePaRank. Given pairwise distances from the model and corresponding contact information from ground-truth we calculate precision and recall curve for a range of thresholds and select one that optimizes F1 score. The resulting threshold allows any residue pair to be classified as ‘in contact’ or ‘not in contact’ based solely on their embedding distance. Finally, we track the precision and recall for contact detection (given the selected threshold) between an antibody and an antigen only (since a high preservation of the antibody structure, i.e. interactions between heavy and light chains are easy to predict). The results are presented in Table 2.

**Table 2.**
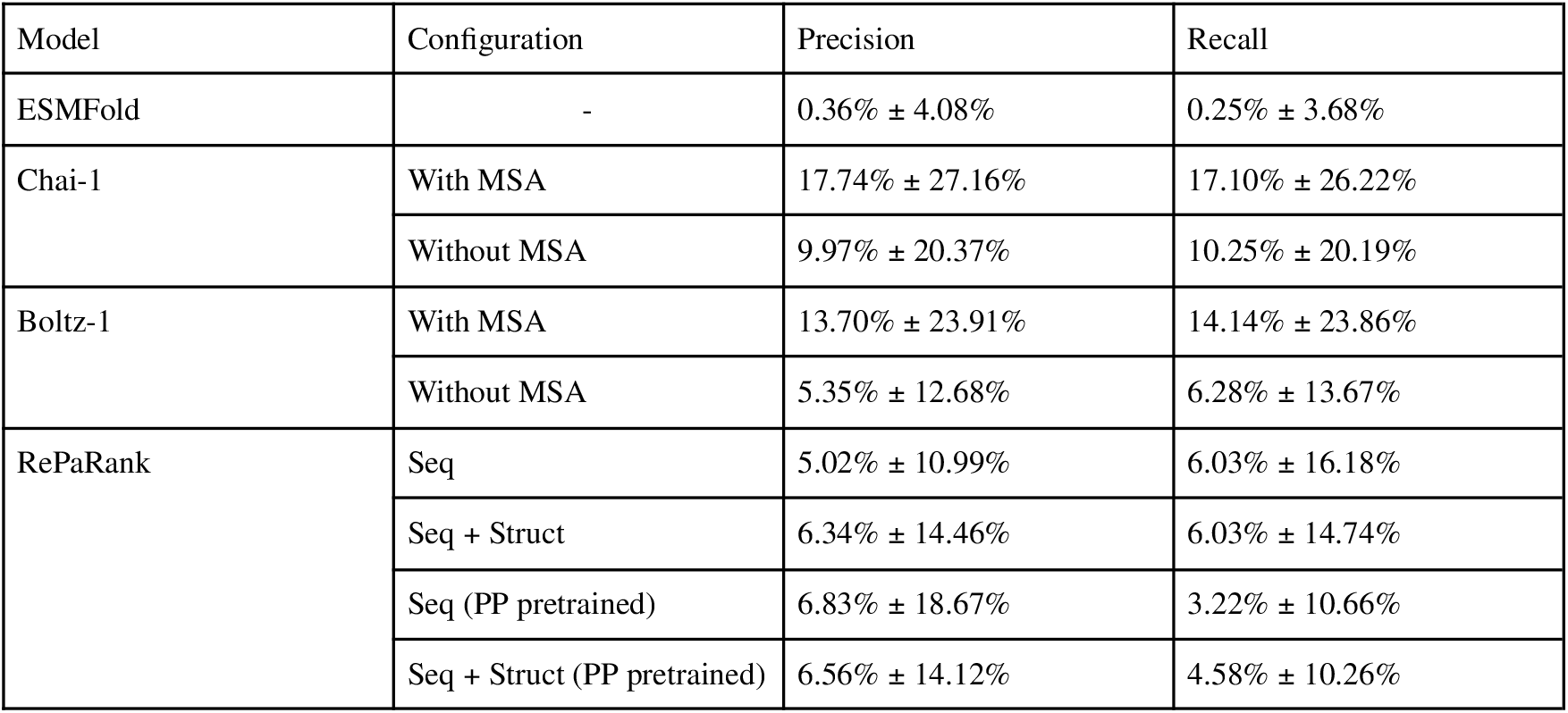
The mean precision and recall (± standard deviation) for identifying contacting residue pairs between antibodies and antigens. A contact is defined by a heavy-atom distance threshold of < 4.5Å. While the MSA-dependent Chai-1 model achieves the highest average scores, a key observation is the high variance across all methods, which suggests inconsistent performance on this task. This binary task is considered less representative of model performance than the Precision AUC ranking metric (see Table 1).

| Model | Configuration | Precision | Recall |
| --- | --- | --- | --- |
| ESMFold | - | 0.36% ± 4.08% | 0.25% ± 3.68% |
| Chai-1 | With MSA | 17.74% ± 27.16% | 17.10% ± 26.22% |
| | Without MSA | 9.97% ± 20.37% | 10.25% ± 20.19% |
| Boltz-1 | With MSA | 13.70% ± 23.91% | 14.14% ± 23.86% |
| | Without MSA | 5.35% ± 12.68% | 6.28% ± 13.67% |
| RePaRank | Seq | 5.02% ± 10.99% | 6.03% ± 16.18% |
| | Seq + Struct | 6.34% ± 14.46% | 6.03% ± 14.74% |
| | Seq (PP pretrained) | 6.83% ± 18.67% | 3.22% ± 10.66% |
| | Seq + Struct (PP pretrained) | 6.56% ± 14.12% | 4.58% ± 10.26% |

The performance analysis across the evaluated protein folding and ranking models reveals a significant performance gap tied to the use of MSA, with Chai-1 (with MSA) and Boltz-1 (with MSA) achieving the highest average precision (17.74% and 13.70%, respectively). However, a critical observation for this dataset is the extreme variance observed across all configurations, where standard deviations frequently exceed the mean values (see Figure 5 for visual interpretation). This statistical volatility suggests that model success is highly target-dependent, with performance likely driven by a subset of high-confidence predictions rather than uniform accuracy across the test set. Consequently, while Chai-1 demonstrates the highest numerical peaks, the overlapping confidence intervals between top-tier models make a definitive ranking of “best” architecture statistically challenging.

**Figure 5.**
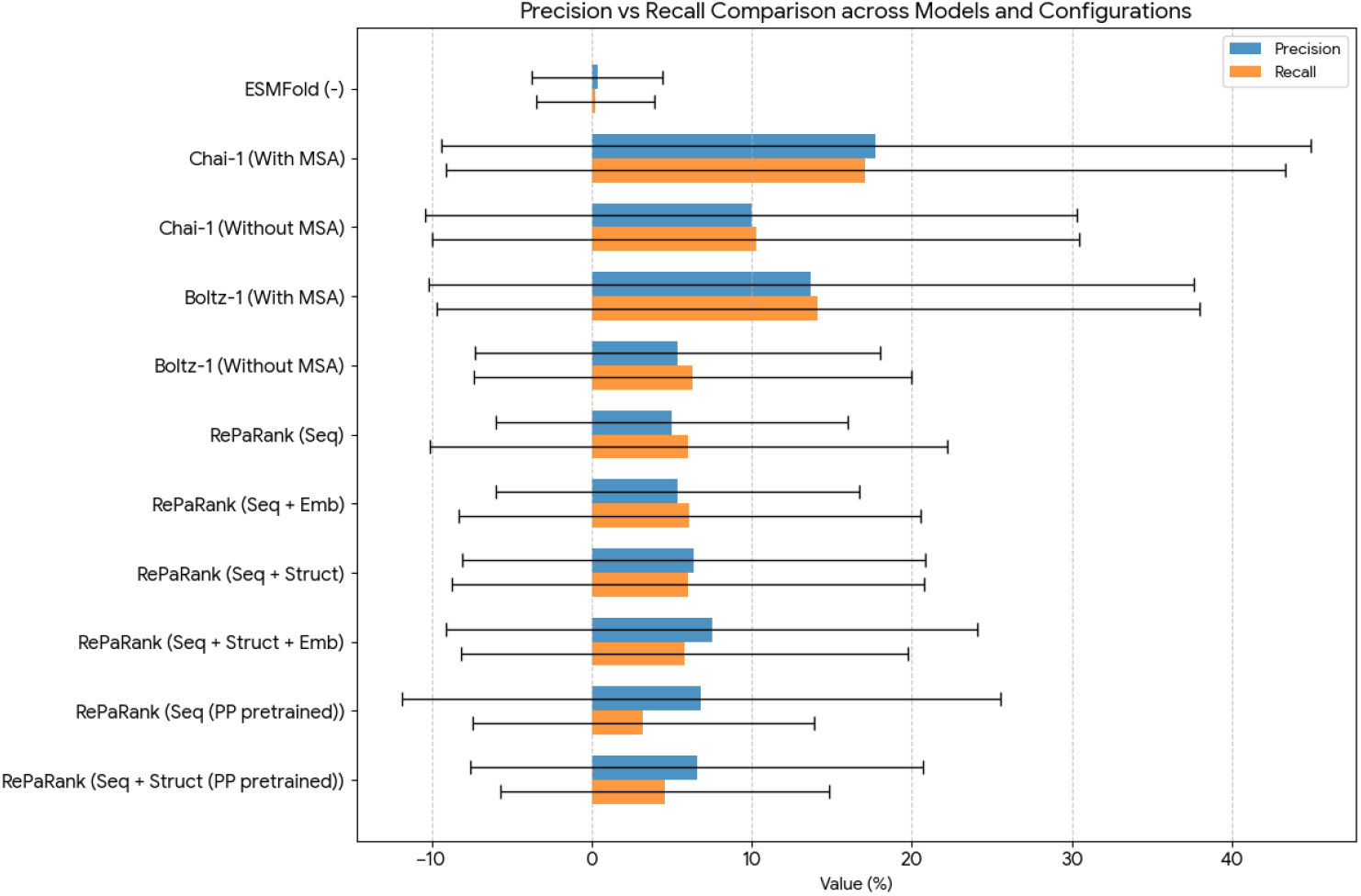
The figure visualizes precision and recall (± standard deviation) from Table 2 for better understanding the instability of the models.

Furthermore, the results highlight a distinct trade-off between model complexity and stability. While MSA-dependent models reach higher performance ceilings, RePaRank configurations show more modest but relatively more consistent metrics. These findings underscore that in the current landscape of structural bioinformatics, the choice of model should be dictated by the specific biological context and the availability of evolutionary information, as no single architecture yet provides reliable, low-variance predictions across all benchmarks.

## 4. Discussion

The primary contribution of this work is the introduction of a lightweight architecture RePaRank designed to capture the intricate geometric properties of protein complexes within a high-dimensional representation space. Unlike existing heavy-parameter models, our approach achieves efficiency without sacrificing the ability to model the subtle spatial dependencies inherent in protein-protein interfaces. To ensure the reliability of our findings, we grounded our evaluation in a methodology centered on Precision AUC. This metric is particularly robust for assessing structure-based prediction models in the context of protein-protein interactions, as it prioritizes the identification of true positive interface residues while effectively handling the class imbalance typical of surface-level binding tasks. By combining geometric sophistication with a rigorous, precision-focused evaluation, we demonstrate that model performance in epitope and paratope prediction is driven more by effective structural representation than by sheer parameter volume.

Traditional binary contact prediction, which relies on a single, arbitrary distance threshold such as 4.5Å, can be misleading. As evidenced by our results in Table 2, this binary task leads to outputs with extremely high variance for all tested models, suggesting performance is inconsistent and unreliable across different complexes. In contrast, the Precision AUC methodology evaluates the model’s entire ranking of potential interactions. This provides a more stable, comprehensive, and practically relevant assessment of a model’s ability to prioritize the most likely interaction sites.

Assessed through this robust framework, RePaRank demonstrates highly competitive performance, given its lightweight implementation. For **paratope** prediction (Table 1; columns *Paratope-H* and *Paratope-L*), RePaRank’s various configurations provide better quality predictions in terms of Precision AUC than other benchmarked models (including Chai-1 and Boltz-1). In the case of **epitope** prediction (Table 1; column *Epitope*), while the MSA-dependent version of Chai-1 achieves the highest absolute score, RePaRank shows a clear performance advantage over the benchmark models when they are used in a self-contained, MSA-free setting. The lack of performance gain from MSA in paratope prediction is consistent with the biology of antibodies; since they do not undergo the long-term co-evolutionary processes typically captured by MSA protocols, traditional evolutionary signals remain uninformative for binding site identification. In contrast, the positive impact of MSA on epitope prediction remains intriguing. Given that epitopes are often perceived as transient surfaces rather than conserved structural motifs, the link between co-evolutionary signals and “epitope-ness” is not immediately clear. This phenomenon suggests a potential contradiction with the standard assumption that epitope recognition is independent of deep evolutionary conservation, warranting further investigation into the structural signals captured by these alignments.

The success of our model can be attributed to its fundamental design: reframing the problem from 3D coordinate prediction to proximity ranking in an N-dimensional embedding space. This approach bypasses the need to explicitly model complex biophysics, simplifying the learning task. This design philosophy results in significant practical advantages. With just 39.4 million parameters, the model is computationally efficient and accessible for training and inference. Furthermore, its independence from external Multiple Sequence Alignment (MSA) databases simplifies the inference pipeline and makes it a more practical tool for high-throughput analysis compared to models that rely on such dependencies.

Despite these strong results, this study has limitations. The top performance of the MSA-dependent Chai-1 on epitope prediction underscores the continued power of deep evolutionary information. It suggests that the processed biological data likely contain a wealth of latent features that remain underutilized by single-sequence models. When successfully paired with external information sources, such as MSA or structural databases, these hidden features can be leveraged to significantly improve performance in high-complexity tasks like protein structure prediction and functional site mapping.

Furthermore, the existence of these intricate, multi-level dependencies suggests that biological data may require more than just increased scale. If the underlying logic of protein interactions involves complex, rule-based constraints alongside statistical patterns, these relationships deserve individual, specialized modeling. Approaches such as neurosymbolic AI, which combine the pattern-recognition strengths of neural networks with the formal reasoning and interpretability of symbolic logic, may be uniquely suited to disentangling these sophisticated biological signals.

Looking forward, several avenues for future work are apparent. First, RePaRank was trained on a relatively small dataset; a significant next step is to scale up the pre-training with a much larger and more diverse structural dataset, including single chains and large multi-protein complexes, to learn more universal geometric representations. Second, since RePaRank was designed for proximity ranking, future work could focus on developing specialized modules that use its embeddings as input for downstream tasks, such as more sophisticated binary contact prediction or guiding protein-protein docking. Finally, the performance gains observed when RePaRank incorporates additional data sources suggest a clear path toward developing specialized components dedicated to the integration of external information.

In conclusion, RePaRank, evaluated through the robust Precision AUC framework, demonstrates that a specialized and computationally efficient model can achieve highly competitive performance against larger, more complex architectures. It offers a practical, accessible, and powerful tool for the immunoinformatics community.

## Supporting information

LICENSE

## Acknowledgements

This research was conducted as a part of the project “Development of the Specitope technology for bioinformatics-based prediction of B-cell-specific epitopes, enabling the ‘in silico’ identification of candidates for therapeutic antibodies.” (Grant No. FEMP.01.01-IP.01-0319/23-00; European Funds).

## Footnotes

1 A per-chain 3D structure was centered, randomly rotated and shifted at the atomic level to mimic independence in a complex.

