## Supplementary material for "RePaRank: An Efficient Architecture for Antibody-Antigen Interface Prediction by Proximity Ranking": LICENSE

Attribution-NonCommercial 4.0 International

=======================================================================

Creative Commons Corporation ("Creative Commons") is not a law firm and
does not provide legal services or legal advice. Distribution of
Creative Commons public licenses does not create a lawyer-client or
other relationship. Creative Commons makes its licenses and related
information available on an "as-is" basis. Creative Commons gives no
warranties regarding its licenses, any material licensed under their
terms and conditions, or any related information. Creative Commons
disclaims all liability for damages resulting from their use to the
fullest extent possible.

Using Creative Commons Public Licenses

Creative Commons public licenses provide a standard set of terms and
conditions that creators and other rights holders may use to share
original works of authorship and other material subject to copyright
and certain other rights specified in the public license below. The
following considerations are for informational purposes only, are not
exhaustive, and do not form part of our licenses.

     Considerations for licensors: Our public licenses are
     intended for use by those authorized to give the public
     permission to use material in ways otherwise restricted by
     copyright and certain other rights. Our licenses are
     irrevocable. Licensors should read and understand the terms
     and conditions of the license they choose before applying it.
     Licensors should also secure all rights necessary before
     applying our licenses so that the public can reuse the
     material as expected. Licensors should clearly mark any
     material not subject to the license. This includes other CC-
     licensed material, or material used under an exception or
     limitation to copyright. More considerations for licensors:
    wiki.creativecommons.org/Considerations_for_licensors

     Considerations for the public: By using one of our public
     licenses, a licensor grants the public permission to use the
     licensed material under specified terms and conditions. If
     the licensor's permission is not necessary for any reason--for
     example, because of any applicable exception or limitation to
     copyright--then that use is not regulated by the license. Our
     licenses grant only permissions under copyright and certain
     other rights that a licensor has authority to grant. Use of
     the licensed material may still be restricted for other
     reasons, including because others have copyright or other
     rights in the material. A licensor may make special requests,
     such as asking that all changes be marked or described.
     Although not required by our licenses, you are encouraged to
     respect those requests where reasonable. More considerations
     for the public:
    wiki.creativecommons.org/Considerations_for_licensees

=======================================================================

Creative Commons Attribution-NonCommercial 4.0 International Public
License

By exercising the Licensed Rights (defined below), You accept and agree
to be bound by the terms and conditions of this Creative Commons
Attribution-NonCommercial 4.0 International Public License ("Public
License"). To the extent this Public License may be interpreted as a
contract, You are granted the Licensed Rights in consideration of Your
acceptance of these terms and conditions, and the Licensor grants You
such rights in consideration of benefits the Licensor receives from
making the Licensed Material available under these terms and
conditions.


Section 1 -- Definitions.

  a. Adapted Material means material subject to Copyright and Similar
     Rights that is derived from or based upon the Licensed Material
     and in which the Licensed Material is translated, altered,
     arranged, transformed, or otherwise modified in a manner requiring
     permission under the Copyright and Similar Rights held by the
     Licensor. For purposes of this Public License, where the Licensed
     Material is a musical work, performance, or sound recording,
     Adapted Material is always produced where the Licensed Material is
     synched in timed relation with a moving image.

  b. Adapter's License means the license You apply to Your Copyright
     and Similar Rights in Your contributions to Adapted Material in
     accordance with the terms and conditions of this Public License.

  c. Copyright and Similar Rights means copyright and/or similar rights
     closely related to copyright including, without limitation,
     performance, broadcast, sound recording, and Sui Generis Database
     Rights, without regard to how the rights are labeled or
     categorized. For purposes of this Public License, the rights
     specified in Section 2(b)(1)-(2) are not Copyright and Similar
     Rights.
  d. Effective Technological Measures means those measures that, in the
     absence of proper authority, may not be circumvented under laws
     fulfilling obligations under Article 11 of the WIPO Copyright
     Treaty adopted on December 20, 1996, and/or similar international
     agreements.

  e. Exceptions and Limitations means fair use, fair dealing, and/or
     any other exception or limitation to Copyright and Similar Rights
     that applies to Your use of the Licensed Material.

  f. Licensed Material means the artistic or literary work, database,
     or other material to which the Licensor applied this Public
     License.

  g. Licensed Rights means the rights granted to You subject to the
     terms and conditions of this Public License, which are limited to
     all Copyright and Similar Rights that apply to Your use of the
     Licensed Material and that the Licensor has authority to license.

  h. Licensor means the individual(s) or entity(ies) granting rights
     under this Public License.

  i. NonCommercial means not primarily intended for or directed towards
     commercial advantage or monetary compensation. For purposes of
     this Public License, the exchange of the Licensed Material for
     other material subject to Copyright and Similar Rights by digital
     file-sharing or similar means is NonCommercial provided there is
     no payment of monetary compensation in connection with the
     exchange.

  j. Share means to provide material to the public by any means or
     process that requires permission under the Licensed Rights, such
     as reproduction, public display, public performance, distribution,
     dissemination, communication, or importation, and to make material
     available to the public including in ways that members of the
     public may access the material from a place and at a time
     individually chosen by them.

  k. Sui Generis Database Rights means rights other than copyright
     resulting from Directive 96/9/EC of the European Parliament and of
     the Council of 11 March 1996 on the legal protection of databases,
     as amended and/or succeeded, as well as other essentially
     equivalent rights anywhere in the world.

  l. You means the individual or entity exercising the Licensed Rights
     under this Public License. Your has a corresponding meaning.


Section 2 -- Scope.

  a. License grant.

       1. Subject to the terms and conditions of this Public License,
          the Licensor hereby grants You a worldwide, royalty-free,
          non-sublicensable, non-exclusive, irrevocable license to
          exercise the Licensed Rights in the Licensed Material to:

            a. reproduce and Share the Licensed Material, in whole or
               in part, for NonCommercial purposes only; and

            b. produce, reproduce, and Share Adapted Material for
               NonCommercial purposes only.

       2. Exceptions and Limitations. For the avoidance of doubt, where
          Exceptions and Limitations apply to Your use, this Public
          License does not apply, and You do not need to comply with
          its terms and conditions.

       3. Term. The term of this Public License is specified in Section
          6(a).

       4. Media and formats; technical modifications allowed. The
          Licensor authorizes You to exercise the Licensed Rights in
          all media and formats whether now known or hereafter created,
          and to make technical modifications necessary to do so. The
          Licensor waives and/or agrees not to assert any right or
          authority to forbid You from making technical modifications
          necessary to exercise the Licensed Rights, including
          technical modifications necessary to circumvent Effective
          Technological Measures. For purposes of this Public License,
          simply making modifications authorized by this Section 2(a)
          (4) never produces Adapted Material.

       5. Downstream recipients.

            a. Offer from the Licensor -- Licensed Material. Every
               recipient of the Licensed Material automatically
               receives an offer from the Licensor to exercise the
               Licensed Rights under the terms and conditions of this
               Public License.

            b. No downstream restrictions. You may not offer or impose
               any additional or different terms or conditions on, or
               apply any Effective Technological Measures to, the
               Licensed Material if doing so restricts exercise of the
               Licensed Rights by any recipient of the Licensed
               Material.

       6. No endorsement. Nothing in this Public License constitutes or
          may be construed as permission to assert or imply that You
          are, or that Your use of the Licensed Material is, connected
          with, or sponsored, endorsed, or granted official status by,
          the Licensor or others designated to receive attribution as
          provided in Section 3(a)(1)(A)(i).

  b. Other rights.

       1. Moral rights, such as the right of integrity, are not
          licensed under this Public License, nor are publicity,
          privacy, and/or other similar personality rights; however, to
          the extent possible, the Licensor waives and/or agrees not to
          assert any such rights held by the Licensor to the limited
          extent necessary to allow You to exercise the Licensed
          Rights, but not otherwise.

       2. Patent and trademark rights are not licensed under this
          Public License.

       3. To the extent possible, the Licensor waives any right to
          collect royalties from You for the exercise of the Licensed
          Rights, whether directly or through a collecting society
          under any voluntary or waivable statutory or compulsory
          licensing scheme. In all other cases the Licensor expressly
          reserves any right to collect such royalties, including when
          the Licensed Material is used other than for NonCommercial
          purposes.


Section 3 -- License Conditions.

Your exercise of the Licensed Rights is expressly made subject to the
following conditions.

  a. Attribution.

       1. If You Share the Licensed Material (including in modified
          form), You must:

            a. retain the following if it is supplied by the Licensor
               with the Licensed Material:

                 i. identification of the creator(s) of the Licensed
                    Material and any others designated to receive
                    attribution, in any reasonable manner requested by
                    the Licensor (including by pseudonym if
                    designated);

                ii. a copyright notice;

               iii. a notice that refers to this Public License;

                iv. a notice that refers to the disclaimer of
                    warranties;

                 v. a URI or hyperlink to the Licensed Material to the
                    extent reasonably practicable;

            b. indicate if You modified the Licensed Material and
               retain an indication of any previous modifications; and

            c. indicate the Licensed Material is licensed under this
               Public License, and include the text of, or the URI or
               hyperlink to, this Public License.

       2. You may satisfy the conditions in Section 3(a)(1) in any
          reasonable manner based on the medium, means, and context in
          which You Share the Licensed Material. For example, it may be
          reasonable to satisfy the conditions by providing a URI or
          hyperlink to a resource that includes the required
          information.

       3. If requested by the Licensor, You must remove any of the
          information required by Section 3(a)(1)(A) to the extent
          reasonably practicable.

       4. If You Share Adapted Material You produce, the Adapter's
          License You apply must not prevent recipients of the Adapted
          Material from complying with this Public License.


Section 4 -- Sui Generis Database Rights.

Where the Licensed Rights include Sui Generis Database Rights that
apply to Your use of the Licensed Material:

  a. for the avoidance of doubt, Section 2(a)(1) grants You the right
     to extract, reuse, reproduce, and Share all or a substantial
     portion of the contents of the database for NonCommercial purposes
     only;

  b. if You include all or a substantial portion of the database
     contents in a database in which You have Sui Generis Database
     Rights, then the database in which You have Sui Generis Database
     Rights (but not its individual contents) is Adapted Material; and

  c. You must comply with the conditions in Section 3(a) if You Share
     all or a substantial portion of the contents of the database.

For the avoidance of doubt, this Section 4 supplements and does not
replace Your obligations under this Public License where the Licensed
Rights include other Copyright and Similar Rights.


Section 5 -- Disclaimer of Warranties and Limitation of Liability.

  a. UNLESS OTHERWISE SEPARATELY UNDERTAKEN BY THE LICENSOR, TO THE
     EXTENT POSSIBLE, THE LICENSOR OFFERS THE LICENSED MATERIAL AS-IS
     AND AS-AVAILABLE, AND MAKES NO REPRESENTATIONS OR WARRANTIES OF
     ANY KIND CONCERNING THE LICENSED MATERIAL, WHETHER EXPRESS,
     IMPLIED, STATUTORY, OR OTHER. THIS INCLUDES, WITHOUT LIMITATION,
     WARRANTIES OF TITLE, MERCHANTABILITY, FITNESS FOR A PARTICULAR
     PURPOSE, NON-INFRINGEMENT, ABSENCE OF LATENT OR OTHER DEFECTS,
     ACCURACY, OR THE PRESENCE OR ABSENCE OF ERRORS, WHETHER OR NOT
     KNOWN OR DISCOVERABLE. WHERE DISCLAIMERS OF WARRANTIES ARE NOT
     ALLOWED IN FULL OR IN PART, THIS DISCLAIMER MAY NOT APPLY TO YOU.

  b. TO THE EXTENT POSSIBLE, IN NO EVENT WILL THE LICENSOR BE LIABLE
     TO YOU ON ANY LEGAL THEORY (INCLUDING, WITHOUT LIMITATION,
     NEGLIGENCE) OR OTHERWISE FOR ANY DIRECT, SPECIAL, INDIRECT,
     INCIDENTAL, CONSEQUENTIAL, PUNITIVE, EXEMPLARY, OR OTHER LOSSES,
     COSTS, EXPENSES, OR DAMAGES ARISING OUT OF THIS PUBLIC LICENSE OR
     USE OF THE LICENSED MATERIAL, EVEN IF THE LICENSOR HAS BEEN
     ADVISED OF THE POSSIBILITY OF SUCH LOSSES, COSTS, EXPENSES, OR
     DAMAGES. WHERE A LIMITATION OF LIABILITY IS NOT ALLOWED IN FULL OR
     IN PART, THIS LIMITATION MAY NOT APPLY TO YOU.

  c. The disclaimer of warranties and limitation of liability provided
     above shall be interpreted in a manner that, to the extent
     possible, most closely approximates an absolute disclaimer and
     waiver of all liability.


Section 6 -- Term and Termination.

  a. This Public License applies for the term of the Copyright and
     Similar Rights licensed here. However, if You fail to comply with
     this Public License, then Your rights under this Public License
     terminate automatically.

  b. Where Your right to use the Licensed Material has terminated under
     Section 6(a), it reinstates:

       1. automatically as of the date the violation is cured, provided
          it is cured within 30 days of Your discovery of the
          violation; or

       2. upon express reinstatement by the Licensor.

     For the avoidance of doubt, this Section 6(b) does not affect any
     right the Licensor may have to seek remedies for Your violations
     of this Public License.

  c. For the avoidance of doubt, the Licensor may also offer the
     Licensed Material under separate terms or conditions or stop
     distributing the Licensed Material at any time; however, doing so
     will not terminate this Public License.

  d. Sections 1, 5, 6, 7, and 8 survive termination of this Public
     License.


Section 7 -- Other Terms and Conditions.

  a. The Licensor shall not be bound by any additional or different
     terms or conditions communicated by You unless expressly agreed.

  b. Any arrangements, understandings, or agreements regarding the
     Licensed Material not stated herein are separate from and
     independent of the terms and conditions of this Public License.


Section 8 -- Interpretation.

  a. For the avoidance of doubt, this Public License does not, and
     shall not be interpreted to, reduce, limit, restrict, or impose
     conditions on any use of the Licensed Material that could lawfully
     be made without permission under this Public License.

  b. To the extent possible, if any provision of this Public License is
     deemed unenforceable, it shall be automatically reformed to the
     minimum extent necessary to make it enforceable. If the provision
     cannot be reformed, it shall be severed from this Public License
     without affecting the enforceability of the remaining terms and
     conditions.

  c. No term or condition of this Public License will be waived and no
     failure to comply consented to unless expressly agreed to by the
     Licensor.

  d. Nothing in this Public License constitutes or may be interpreted
     as a limitation upon, or waiver of, any privileges and immunities
     that apply to the Licensor or You, including from the legal
     processes of any jurisdiction or authority.

=======================================================================

Creative Commons is not a party to its public
licenses. Notwithstanding, Creative Commons may elect to apply one of
its public licenses to material it publishes and in those instances
will be considered the “Licensor.” The text of the Creative Commons
public licenses is dedicated to the public domain under the CC0 Public
Domain Dedication. Except for the limited purpose of indicating that
material is shared under a Creative Commons public license or as
otherwise permitted by the Creative Commons policies published at
creativecommons.org/policies, Creative Commons does not authorize the
use of the trademark "Creative Commons" or any other trademark or logo
of Creative Commons without its prior written consent including,
without limitation, in connection with any unauthorized modifications
to any of its public licenses or any other arrangements,
understandings, or agreements concerning use of licensed material. For
the avoidance of doubt, this paragraph does not form part of the
public licenses.

Creative Commons may be contacted at creativecommons.org.
